# Niemann-Pick C-like endo-lysosomal dysfunction in DHDDS patient cells, a congenital disorder of glycosylation, can be treated with miglustat

**DOI:** 10.1101/2025.01.08.631939

**Authors:** Hannah L. Best, Sophie R. Cook, Helen Waller-Evans, Emyr Lloyd-Evans

## Abstract

DHDDS (dehydrodolichol diphosphate synthetase) and NgBR (Nogo-B Receptor) collectively form an enzymatic complex important for the synthesis of dolichol – a key component of protein N-glycosylation. Mutations in *DHDDS* and the gene encoding NgBR (*NUS1)* are associated with neurodevelopmental disorders that clinically present with epilepsy, motor impairments, and developmental delay. Previous work has demonstrated both DHDDS and NgBR can also interact with NPC2 (Niemann-Pick C (NPC) type 2) – a protein which functions to traffic cholesterol out of the lysosome and, when mutated, can cause a lysosomal storage disorder (NPC disease) characterised by an accumulation of cholesterol and glycosphingolipids. Abnormal cholesterol accumulation has also been reported in cells from individuals and animal models with mutations in NUS1, and suspected lipid storage has been shown in biopsies from individuals with mutations in DHDDS. Our findings provide further evidence for overlap between NPC2 and DHDDS disorders, showing that DHDDS patient fibroblasts have increased lysosomal volume, store cholesterol and ganglioside GM1, and have altered lysosomal Ca^2+^ homeostasis. Treatment of DHDDS cells - with the approved NPC small molecule therapy miglustat - improves these disease-associated phenotypes, identifying a possible therapeutic option for DHDDS patients. These data suggest that treatment options currently approved for NPC disease may be translatable to DHDDS/NUS1 patients.

## Introduction

*De novo* mutations in the genes *DHDDS* encoding dehydrodolichol diphosphate synthetase (DHDDS) and *NUS1* (nuclear undecaprenyl pyrophosphate synthase 1) encoding the Nogo-B receptor (NgBR), cause neurodevelopmental syndromes largely characterised by epilepsy and movement abnormalities including mild ataxia, myoclonus, and dystonia [1–3]. Although most cases of DHDDS and NUS1 syndrome are heterozygous, some instances of homozygosity have also been reported whilst compound heterozygosity is rare and associated with a more severe disease presentation [4–6]. The DHDDS and NgBR protein products act together as subunits of the cis-prenyltransferase (cis-PTase) enzyme acting within the mevalonate pathway – a metabolic pathway responsible for the production of isoprenoids and sterols [7–9]. *DHDDS* encodes the catalytic subunit responsible for synthesis of dolichol monophosphate, a lipid that is in turn important for protein N-glycosylation in the endoplasmic reticulum (ER) [10,11]. For these reasons DHDDS and NUS1 are classed as congenital disorders of glycosylation (CDGs), a family of diseases caused by mutations in genes associated with N-glycan biosynthesis [12–14]. Surprisingly, whilst homozygosity for NUS1 or DHDDS is associated with phenotypes resembling a congenital disorder of glycosylation [15,16], heterozygous DHDDS patients do not appear to present with any significant serum glycoprotein hypoglycosylation and, in contrast to other CDGs, urinary dolichol D18/D19 ratios and serum transferrin N-glycosylation profiles are normal, suggesting that other factors may contribute to the pathogenesis of this group of disorder [1,17,18].

It has been shown by several groups that both DHDDS and NgBR bind to the lysosomal cholesterol transport protein NPC2. This specific interaction between DHDDS and NPC2 was first identified from a yeast two-hybrid screen and confirmed by co-IP [19]. An interaction between NgBR and NPC2 was shown in 2009 [20]. This later study also provided evidence that the interaction stabilised NPC2, as knock down of NgBR resulted in reduced NPC2 protein levels whilst increased expression of NgBR resulted in retention of NPC2 in the ER and prelysosomal compartments. Finally, the presence of free cholesterol accumulation, similar to that observed in cells from individuals with mutations in NPC2, was observed in a human HUVEC cell line following knock down of NgBR and in murine embryonic fibroblasts heterozygous for NUS1 [20]. Furthermore, the accumulation of cholesterol has subsequently been reported in NUS1 patient fibroblasts, nus1 morphant zebrafish [21], and in the drosophila model [22], whilst altered lysosomes containing intramembraneous lipid whorls have been noted in DHDDS patient biopsies suggestive of glycosphingolipid (GSL) storage [1].

GSL accumulation is a key hallmark of Niemann-Pick C (NPC) disease, an inherited lysosomal disease caused by mutation and loss of function of the lysosomal cholesterol and lipid handling NPC1 and NPC2 proteins [23]. Despite evidence that disruption of the functional interaction between the subunits of cis-PTase and NPC2 results in NPC2 mis-localisation to the ER, no study has yet confirmed the presence of other hallmarks of NPC disease cellular pathogenesis in DHDDS or NUS1 cells, namely lysosomal storage of other lipids including GSL and lyso-(bis)phosphatidic acid (LBPA), endosomal transport-autophagy abnormalities and lysosomal Ca^2+^ signalling defects [23–25]. As there are now multiple approved disease-modifying therapies for NPC disease, miglustat [26], arimoclomol [27] and N-acetyl-L-leucine [28], demonstrating the presence of these phenotypes in DHDDS cells, where NPC2 function may be impaired, is critical for determining whether these NPC therapies could be repurposed for DHDDS/NUS1 syndrome where there is currently no disease modifying therapy.

## Materials and Methods

### Cell maintenance and drug treatment

Apparently healthy (control) human (GM05399) and NPC2 patient (GM18455) fibroblast cell lines were obtained from the Coriell Cell Repository (New Jersey, USA), DHDDS patient fibroblasts (**Table 1**) were from Dr. Frances Elmslie (St. George’s University Hospital, London, UK) and Dr. Eva Morava (Icahn School of Medicine at Mount Sinai). Fibroblasts were grown as monolayers in Dulbecco’s modified Eagle’s high glucose medium (DMEM, Gibco) supplemented with 10% foetal bovine serum (FBS, PAN biotech) and 1% L-glutamine (complete medium), cultures were maintained in a humidified 5% CO_2_ incubator at 37°C. Miglustat (Toronto Research Chemicals, M344225) was dissolved in mqH_2_0 to generate a 100 mM stock solution, which was stored at –20 °C. For miglustat treatment, stock was diluted in complete media to a final concentration of 50 μM. Fresh miglustat-containing media was added every third day for 14 days prior to live staining, fixation and imaging. For all experiments fibroblasts were passage matched with equal cell numbers seeded, cells were plated on Ibidi chamberslides for Ca^2+^ imaging or Perkin Elmer 96-well PhenoPlates for fluorescent stain imaging.

**Table 1.** Human fibroblast cells used in this study.

| Cell Line | Mutation | Source | Reported DHDDS/NUS1 activity |
| --- | --- | --- | --- |
| WT | Apparently healthy control | Coriell GM01652, Female, 11 yr | Not reported, presumed normal |
| NPC2 | Compound heterozygous c.58G>T, p.Glu20Ter c.140G>T, p.Cys47Phe | Coriell GM18455, Male, no reported age | Not reported, presumed normal |
| DHDDS P1 | c.614 G>A p.Arg205Gln | St. George's University Hospital, Male, 11 yr | ~15-fold lower (at detection limit) [1] |
| DHDDS P2 | c.110 G>A p.Arg37His | St. George's University Hospital, Female, 14 yr | ~ 5-fold decrease [29]<br>~15-fold lower (at detection limit) [1] |
| DHDDS P3 | c.110 G>A p.Arg37His | St. George's University Hospital, Male, 15 yr | ~ 5-fold decrease [29]<br>~15-fold lower (at detection limit) [1] |
| DHDDS P4 | c.614 G>A p.Arg205Gln | Mayo Clinic, Female, 13 yr | ~15-fold lower (at detection limit) [1] |

### Determination of lysosomal Ca^2+^

To determine lysosomal Ca^2+^ levels, cells were first loaded with the membrane permeant ratiometric Ca^2+^ binding dye Fura-2,AM (Abcam, ab120873). Cells were washed and incubated for 45 minutes at room temperature in DMEM with 10% FBS, 1% BSA (Sigma-Aldrich), 0.025% Pluronic F127 (Sigma-Aldrich) and 5μM Fura-2,AM. Following incubation, cells were washed once and incubated for a further 10 minutes at room temperature in DMEM with 10% FBS to allow de-esterification of the dye. Cells were then washed and imaged in Hank’s balanced salt solution supplemented with 5mM HEPES (pH7.4) and 1 mM MgCl_2_ and 1 μM CaCl_2_. Imaging was performed on a Zeiss Axio Observer A1 microscope with Colibri LED illumination and high speed Axiocam Mrm camera. Fura-2,AM was excited at 360 and 380nm LED wavelengths with emission monitored at 510nm, regions of interest were placed over whole cells and videos were recorded using Axiovision 4.8.2 imaging software at intervals of 1-2s following additions. To release lysosomal Ca^2+^, cells were first incubated with the Ca^2+^ ionophore ionomycin (Sigma-Aldrich, I9657), which does not impact lysosomes but causes other membranes to be permeant to Ca^2+^, followed by the cathepsin C substrate Gly-Phe-β-naphthylamide (GPN, Abcam ab145914) whose products induce perforation of the lysosomal membrane triggering release of the intralumenal Ca^2+^. Data was analysed in Microsoft Excel and Graphpad Prism (v10.1.0), Ca^2+^ traces are shown as the ratio of emission at 360/380nm.

### Lysosomal staining with Lysotracker red

Lysosomal density and distribution were determined by live imaging using the lysosomal dye LysoTracker Red DND-99 (Invitrogen, L7528). Cells were loaded at room temperature with 200nM Lysotracker red in complete media for 10 minutes. Following loading, cells were washed once in DPBS, counterstained with the nuclear marker Hoechst 33341 (0.4μg/ml in DPBS, Invitrogen) for 10 minutes at room temperature, washed a further two times in DPBS and imaged live using an Operetta high content imaging system (Perkin Elmer).

### Autophagic vacuole staining with CytoID

The cellular content of autophagic vacuoles, under rest, was determined using the fluorescent small molecule marker CYTO-ID from the CYTO-ID detection kit (ENZO, ENZ-51031) in live cells. Cells were incubated with CYTO-ID according to manufacturers instructions. Briefly, CYTO-ID was diluted 1:1000 in complete medium and cells were incubated for 30 minutes at room temperature prior to washing in DPBS and counterstaining with Hoechst as described above. Cells were imaged live an Operetta high content imaging system (Revvity).

### Staining of cellular lipid content

To determine lipid content and localisation we first washed cells in DPBS followed by fixation in 4% paraformaldehyde for 10 minutes at room temperature. Cells were subsequently washed twice with DPBS. To image cholesterol, cells were stained with the autofluorescent polyene antibiotic filipin complex (BF162725, Carbosynth) at a concentration of 187.5μg/ml in DMEM with 10% FBS for 30 minutes in the dark at room temperature. Cells were then washed once with DMEM with 10% FBS and twice with DPBS. Prior to imaging cells were counterstained with DRAQ5 nuclear marker (ab108410, abcam). To image glycosphingolipids cells were stained with FITC labelled Cholera toxin subunit B (CtxB, C1655, Sigma-Aldrich), a toxin that binds to ganglioside GM1. Fixed cells were incubated with 1μg/ml FITC-CtxB overnight in blocking buffer (DPBS supplemented with 0.01% saponin and 1% BSA) prior to 3 x 5min washes in DPBS and counterstaining of nuclei with Hoechst as described above. Cells were imaged in DPBS using an Operetta high content imaging system (Perkin Elmer).

### Immunocytochemistry

For lysosomal staining a LAMP1 antibody was used (Santa Cruz, sc20011, lotk2222, 1:500). Cells were fixed in 4% PFA as described above, and permeabilised with blocking buffer for 2 h at room temperature. Primary antibodies were added in blocking buffer overnight at 4 °C. After three 5 minute DPBS washes, secondary antibodies were added for 1 h at room temperature (AlexFluor 594 ab150116, AlexaFluor 488 ab150077, both Abcam, 1:500). Cells were washed a further three times for 5 minutes in DPBS and counter stained with Hoescht, as described above. For imaging, cells were left in DPBS.

### Imaging and image analysis

For high content imaging, cells were imaged using an Operetta CLS High-Content Analysis System with Harmony software (revvity). For each ‘n’, duplicate wells were imaged, capturing 40 images per well. Image analysis utilised the affiliated Harmony High-content imaging and analysis software which provides routine automated quantification of cellular phenotypes, including spot count, spot area, and colocalization. For confocal imaging the samples were examined using a Zeiss Cell Discoverer 7 equipped with a Zeiss Axiocam 712 monochrome CMOS camera, the LSM900 series confocal scanhead, and Zen software suite. For each ‘n’ duplicate wells were imaged, capturing 10 images per well using laser lines 405nm, 488nm, 561nm.

### Statistics

Three or four independent experiments were performed per stain as stated in the figure legend as ‘n =’. GraphPad Prism (v. 10.1.0) was used to analyse the data. Two-way ANOVA was used to measure a response affected by two factors (genotype and treatment), followed by Dunnett’s multiple comparisons test comparing the untreated apparently healthy control group to all other groups.

## Results

### NPC2 human fibroblasts present with classical NPC disease cellular phenotypes

Whilst phenotypic characterisation of NPC1 mutant human fibroblasts has been reported frequently in the literature [30–33], to the best of our knowledge, beyond analysis of the NPC2 hypomorph mouse model [34], no one has characterised the presence of phenotypes - other than cholesterol storage - in NPC2 patient fibroblasts [35]. Furthermore, no one, to our knowledge, has confirmed that miglustat mediates benefit in any model of NPC2 disease. To generate a point of reference for the phenotypic analysis of the DHDDS patient cells we first determined what phenotypes were present in NPC2 disease. As expected, we observed a clear accumulation of cholesterol (∼12-fold) and ganglioside GM1 (∼3-fold) in punctate peri-nuclear lysosomal structures in NPC2 patient fibroblasts (**Figure 1**). This lipid accumulation was associated with elevated lysotracker fluorescence (∼2.6-fold) and an accumulation of autophagosomes (∼3.8-fold, **Figure 1**). Whilst these phenotypes are observed in many lysosomal diseases, albeit often at lower levels (e.g. cholesterol) [36], one phenotype that is relatively specific for NPC disease is the reduction of lysosomal Ca^2+^ content caused by the lysosomal accumulation of sphingosine [24]. Using GPN to perforate the lysosome in cells pre-clamped with ionomycin, we observed a clear reduction in lysosomal Ca^2+^ content in the NPC2 cells (**Figure S1**). In combination with significant intra-lysosomal accumulation of cholesterol and gangliosides these phenotypes are classical hallmarks of NPC disease and are indeed present in both NPC1 and NPC2 patient cells, as would be expected. Miglustat is an approved therapeutic for NPC disease that works by reducing GSL biosynthesis via inhibition of glucosylceramide synthase. Treatment of NPC2 disease cells with miglustat, at 50 μM for 14 days, was sufficient to improve all the reported phenotypes except for cholesterol storage (**Figure 1**). This is expected, given that miglustat does not alter cholesterol accumulation in NPC1 mutant cells [26].

**Figure 1.**
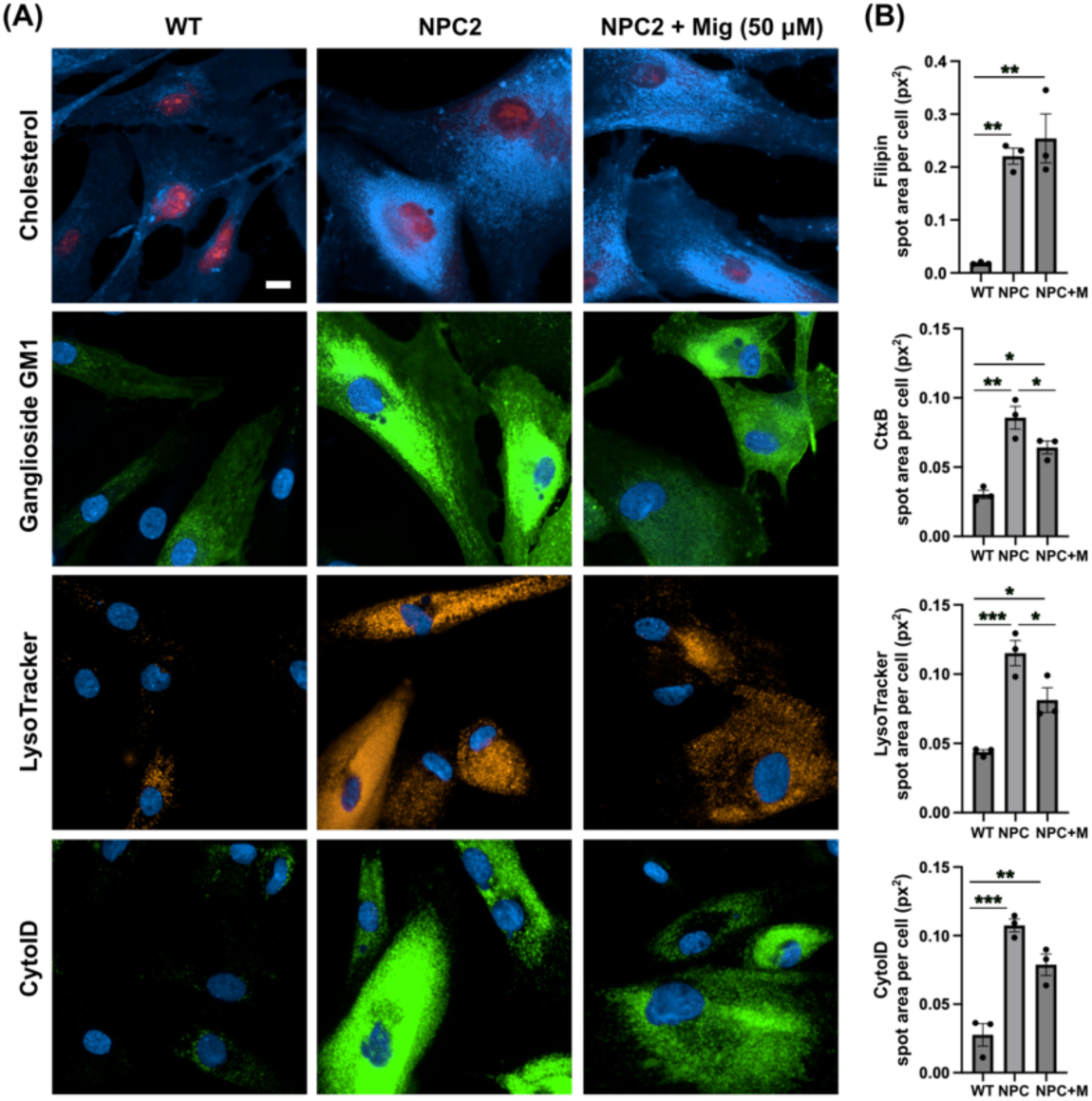
Classical NPC disease phenotypes present in NPC2 human patient fibroblast cells are partially corrected with miglustat. **(A)** Representative images of wild-type (WT) and Niemann Pick C type 2 (NPC2) patient fibroblasts. Fixed cells were stained with filipin to detect cholesterol and FITC-conjugated cholera toxin (CtxB) to detect ganglioside GM1. Live cells were stained with LysoTracker to detect lysosomes and Cyto-ID to detect autophagic vacuoles. Cells are either untreated or treated with 50 μM miglustat (mig, +M) for 14 days. Scale bar = 10 μm. **(B)** Average total spot area per cell for each stain. Data is presented as the mean ± SEM, as determined by ANOVA comparing all groups to the wild-type untreated control (*P ≤ 0.05, **P ≤ 0.01, n = 3).

### DHDDS human fibroblasts cell present with lysosomal lipid storage

For comparative phenotyping assessment in DHDDS patient fibroblasts we used four cell lines, which cover two different mutations (**Table 1**). Whilst cholesterol accumulation has been shown previously across several NUS1 models [21, 22], it has not been fully investigated or confirmed as being lysosomal in DHDDS. Combined filipin and LAMP1 immunofluorescence confirmed a ∼3.4-fold increase in lysosomal cholesterol in all four DHDDS patient cell lines (**Figure 2A,B**). Furthermore, LAMP1 was elevated ∼2.8-fold in all four cell lines, indicating increased number/volume of lysosomes – presumably as a downstream effect of lipid storage (**Figure 2A,C**). Miglustat treatment has no effect on cholesterol accumulation, which is unsurprising given we know it has no beneficial effect in NPC disease models. LAMP1 also remained unchanged with miglustat treatment. Using combined CtxB and LAMP1 staining, we went on to examine the intracellular distribution of a glycosphingolipid, ganglioside GM1 (**Figure 3A**). In the apparently healthy control cells, ganglioside GM1 is present in cellular structures that do not colocalise with the lysosomal LAMP1 marker, whilst in the DHDDS cell lines we see an accumulation of perinuclear punctate LAMP1-positive structures containing ganglioside GM1 (**Figure 3A**). Miglustat treatment significantly reduced the punctate staining of ganglioside GM1 by ∼1.5-fold, confirming that miglustat inhibits GSL synthesis resulting in reduced lysosomal storage in DHDDS patient cells (**Figure 3B**). Whilst miglustat treatment decreased GSL storage, LAMP1 remained unchanged – as quantified in Figure 2 - suggesting lysosomal expression of LAMP1 does not decrease in the same timeframe. After confirmation of lipid storage, we next looked at LysoTracker – a marker of lysosomal volume that serves as a surrogate indicator for lysosomal storage and commonly used as a biomarker for NPC and other lysosomal storage disorders [37] (**Figure 4A**). LysoTracker was significantly elevated ∼1.7-fold in all DHDDS patient lines, confirming the increased lysosomal volume, as expected from previously observed lipid storage. Miglustat treatment significantly reduced LysoTracker area ∼1.4-fold, indicating decreased lysosomal volume (**Figure 4B**). The reduction of LysoTracker in the absence of a LAMP1 reduction suggests lysosomes are reduced in size but are not being cleared over the 14-day treatment window.

**Figure 2.**
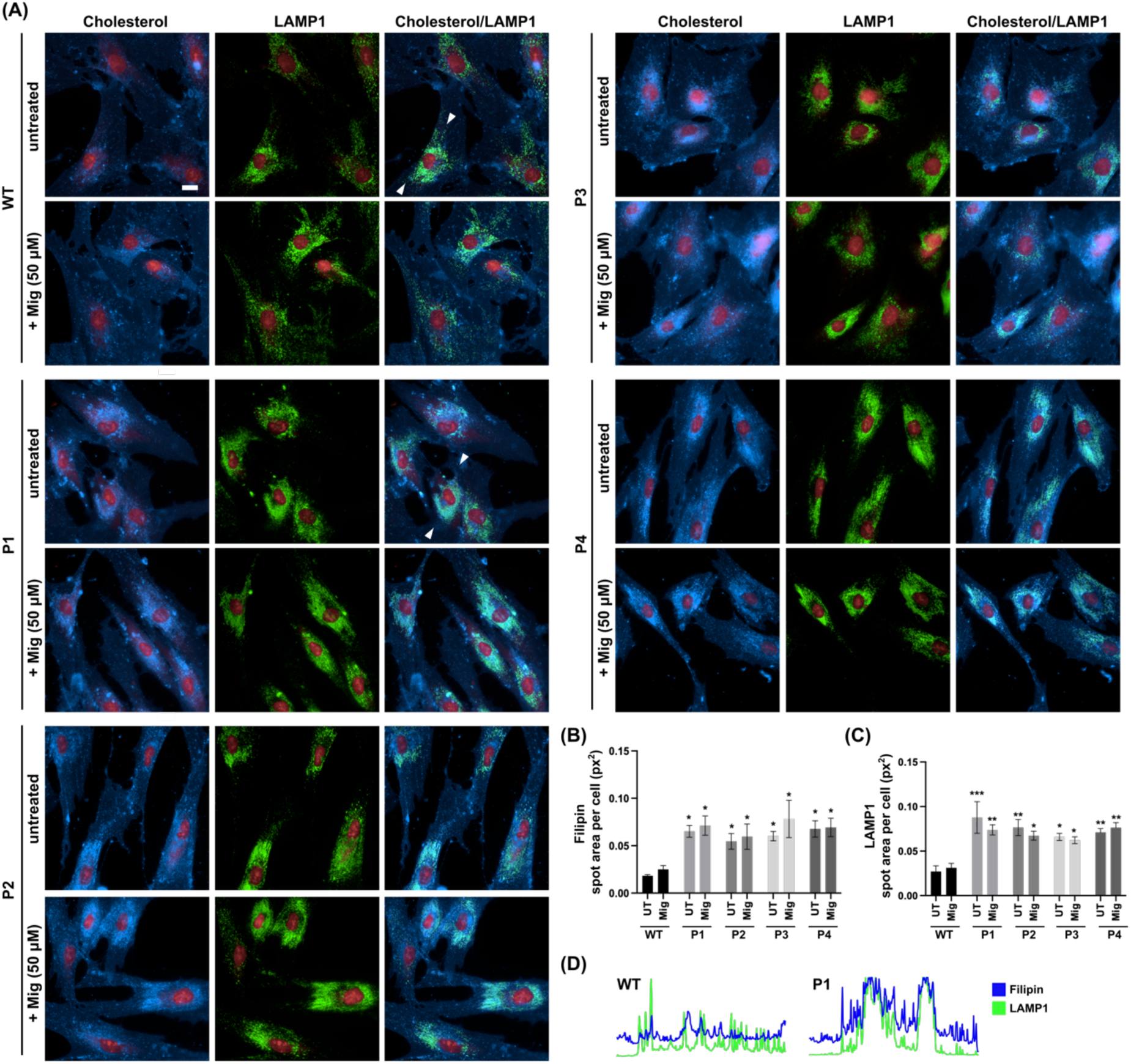
Lysosomal accumulation of cholesterol in DHDDS patient fibroblasts is not altered with miglustat. **(A)** Representative images of wild-type (WT) and DHDDS patient fibroblasts (P1, P2, P3 C P4) fixed and stained with filipin to detect cholesterol and anti-LAMP1 to detect lysosomes. Cells are either untreated or treated with 50 μM miglustat (mig) for 14 days. Representative scale bar = 10 μm. **(B)** Filipin average total spot area per cell. **(C)** LAMP1 average total spot area per cell. Data is presented as the mean ± SEM, as determined by ANOVA comparing all comparing all groups to the wild-type untreated control and comparing DHDDS untreated vs miglustat treated (*P ≤ 0.05, **P ≤ 0.01, ***P ≤ 0.01n = 3 – 4). **(D)** Plots of fluorescent intensity from the regions indicated by the arrows in WT and P1.

**Figure 3.**
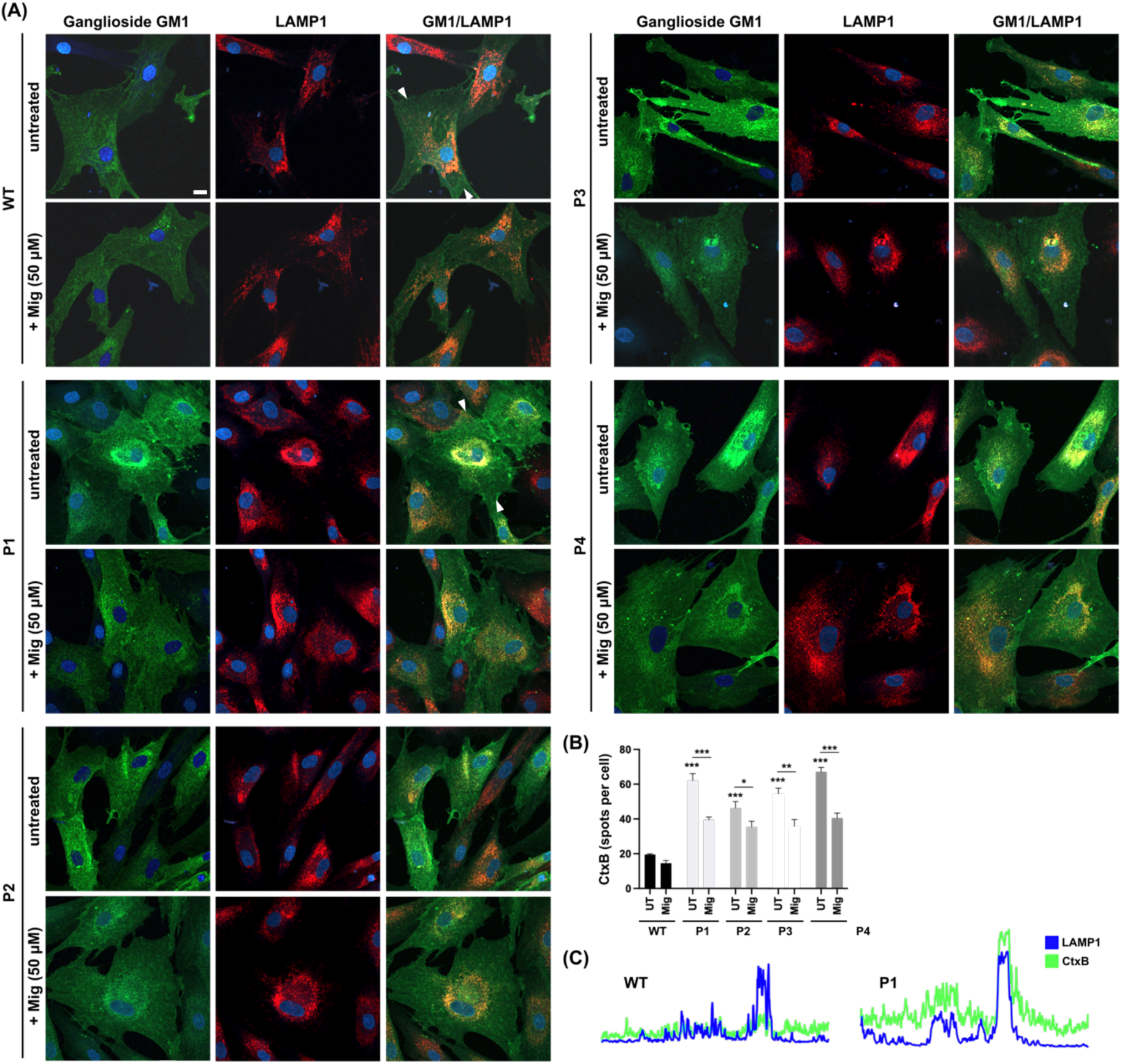
Lysosomal accumulation of ganglioside GM1 in DHDDS patient fibroblasts is reduced with miglustat. **(A)** Representative images of wild-type (WT) and DHDDS patient fibroblasts (P1, P2, P3 C P4) fixed and stained with FITC-conjugated cholera toxin (CtxB) to detect ganglioside GM1 and anti-LAMP1 to detect lysosomes. Cells are either untreated or treated with 50 μM miglustat (Mig) for 14 days. Scale bar = 10 μm. **(B)** CtxB number of spots per cell. Data is presented as the mean ± SEM, as determined by ANOVA comparing all groups to the untreated control and comparing between DHDDS untreated vs miglustat treated (**P ≤ 0.01, ***P ≤ 0.001, n = 3 – 4). **(C)** Plots of fluorescent intensity from the regions indicated by the arrows in WT and P1.

**Figure 4.**
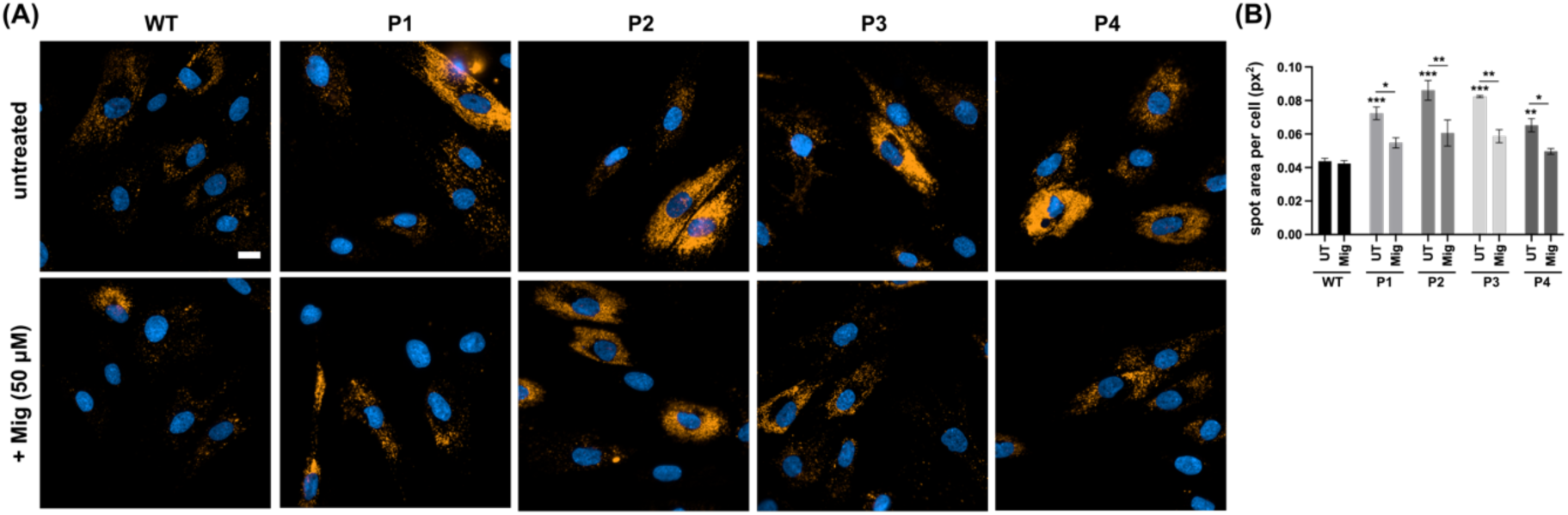
Increased lysosomal volume in DHDDS patient fibroblasts is reduced with miglustat. **(A)** Representative images of wild-type (WT) and DHDDS patient fibroblasts (P1, P2, P3 C P4) live stained with LysoTracker to detect lysosomes. Cells are either untreated or treated with 50 μM miglustat (Mig) for 14 days. Scale bar = 10 μm. **(B)** LysoTracker average spot area per cell. Data is presented as the mean ± SEM, as determined by ANOVA comparing all groups to the wild-type untreated control (*P ≤ 0.05, **P ≤ 0.01, ***P ≤ 0.001, n = 3 – 4).

### DHDDS cells have reduced lysosomal Ca^2+^ content

The storage of lipids, specifically sphingosine, in NPC causes a reduction in lysosomal Ca^2+^ content. To investigate lysosomal Ca^2+^ stores in DHDDS, cells were loaded with the cell permeable Ca^2+^ indicator, Fura2,AM. Cells were first treated with ionomycin to empty non-lysosomal intracellular Ca^2+^ stores, followed by GPN to perforate the lysosomal membrane and glycocalyx, and release lysosomal Ca^2+^ into the cytoplasm (**Figure 5A**) [38, 39]. DHDDS cells had an approx. 50% reduction in lysosomal Ca^2+^ content, which was partially corrected (approx. 30%) with miglustat (**Figure 5B**). Not only does this demonstrate another similarity to NPC but also indicates miglustat could provide a functional benefit in DHDDS fibroblasts.

**Figure 5.**
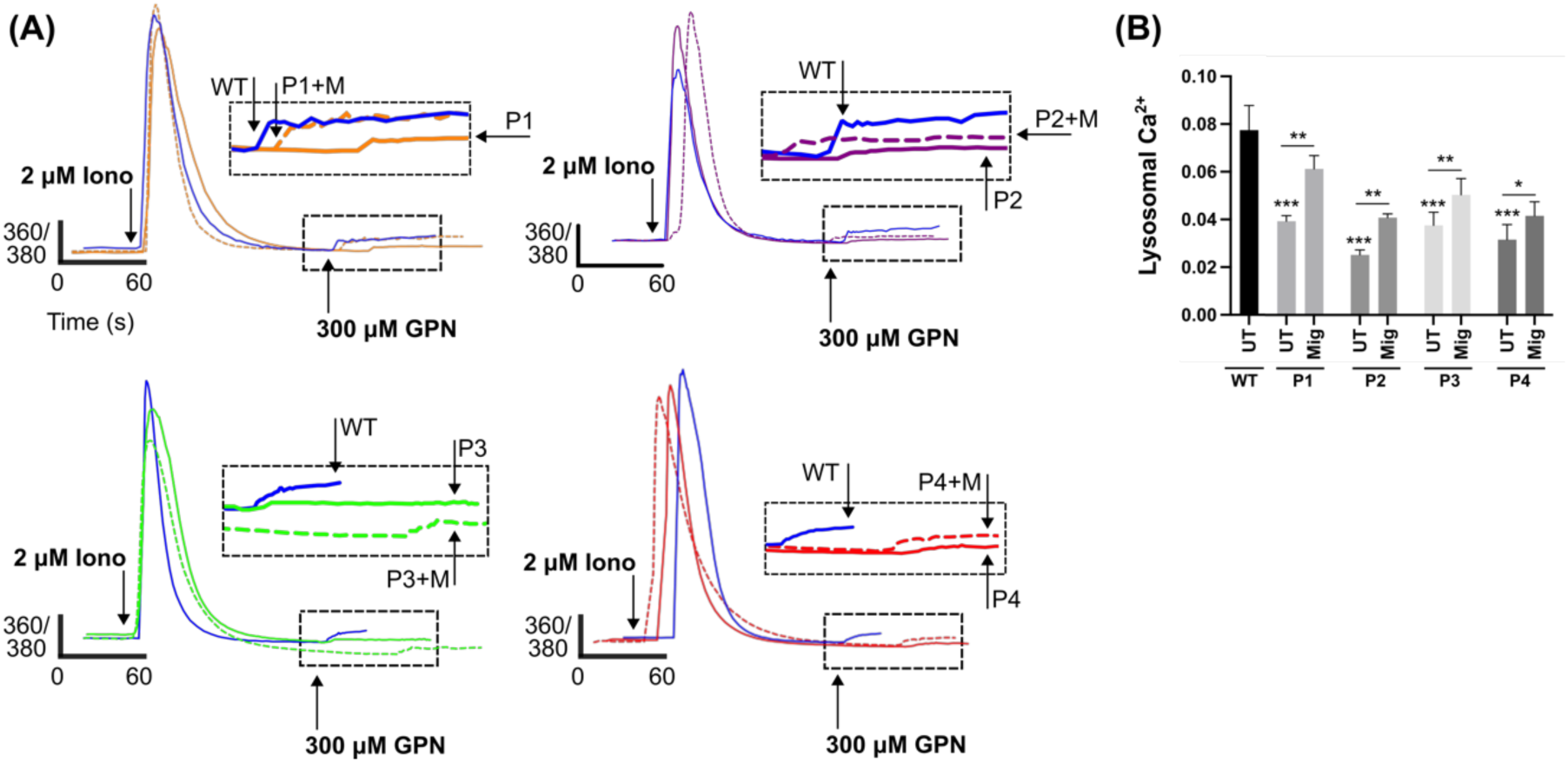
Reduced lysosomal Ca^2+^ in DHDDS patient fibroblasts is partially corrected with miglustat. **(A)** Representative traces of DHDDS patient fibroblasts (P1, P2, P3 C P4) loaded with Fura-2,AM were treated pharmacological agents, ionomycin (iono) and (GPN) to induce Ca^2+^ release. Cells are either untreated or treated with 50 μM miglustat (+M) for 14 days. **(B)** Lysosomal Ca^2+^ release was quantified after GPN addition as indicated and detected by Fura-2,AM (ratiometric measurement at 360 nm and 380 nm, 360/380). Data is presented as the mean ± SEM, as determined by ANOVA comparing DHDDS untreated groups to the wild-type untreated control and comparing DHDDS untreated vs miglustat treated (*P ≤ 0.05, **P ≤ 0.01, ***P ≤ 0.001, n = 4 – 5).

### DHDDS cells have altered autophagic ffux

Finally, we looked at autophagy using the CYTO-ID autophagy probe, another marker which is frequently utilised in lysosomal storage disorders (**Figure 6A**). Autophagic vacuoles were increased ∼2.8-fold across all four patient lines, indicating an upregulation of autophagy or an accumulation and failure to clear autophagic vacuoles. Whilst treatment with miglustat results in a qualitative visible reduction in CYTO-ID intensity between DHDDS cells and apparently healthy controls, there is no significant reduction in spot area (**Figure 6B**). Further analysis did indicate a statistically significant decrease in spot intensity in three patient lines. Further work is needed to determine whether the flux of autophagy is impaired in DHDDS and how this might impact on miglustat treatment.

**Figure 6.**
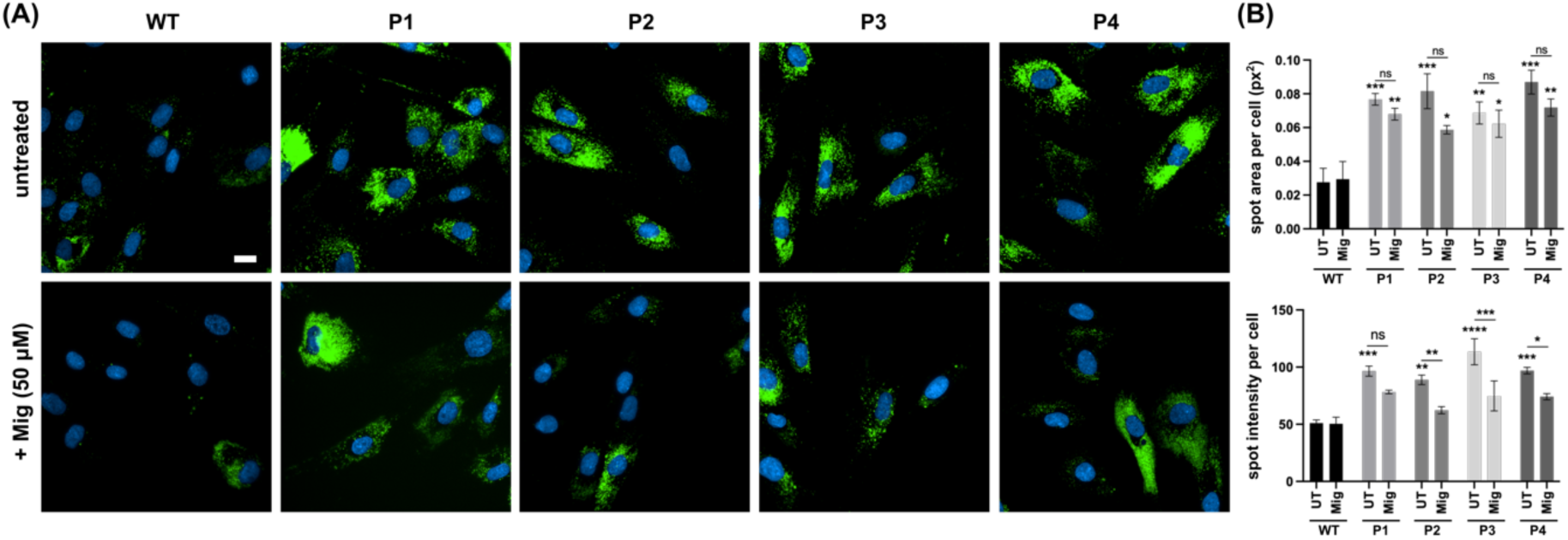
Increased autophagic vacuoles in DHDDS patient fibroblasts. **(A)** Representative images of wild-type (WT) and DHDDS patient fibroblasts (P1, P2, P3 C P4) live stained with Cyto-ID to detect autophagic vacuoles. Cells are either untreated or treated with 50 μM miglustat (mig) for 14 days. Scale bar = 10 μm. **(B)** Cyto-ID average total spot area per cell and spot intensity per cell. Data is presented as the mean ± SEM, as determined by ANOVA comparing all groups to the wild-type untreated control and comparing DHDDS untreated vs miglustat treated (*P ≤ 0.05, **P ≤ 0.01, ***P ≤ 0.001, ***P≤0.0001, n = 3 – 4).

## Discussion

In this report we have demonstrated for the first time, to our knowledge, the presence of a plethora of NPC disease-like phenotypes in DHDDS patient cells. Albeit milder than those we observe in NPC2 loss of function patient cells, all four DHDDS cell lines presented with lysosomal expansion, storage of cholesterol and ganglioside GM1, and increased autophagic vacuoles. Furthermore, we observe a defect in the maintenance of lysosomal Ca^2+^ homeostasis demonstrating a pathogenic change in cell signalling that we know from previous work is a direct consequence of sphingosine storage [24]. Our observations are based on data from four different patient fibroblast cells harbouring two different mutations, located in one of each of the two substrate binding sites present in the cis-PT active site. The functional defect from these mutations is clear, due to both R37 and R205 participating in substrate binding [29]. Both mutations result in a substantial decrease in cis-PT activity to near the lower end of the detectable range (**Table 1**) [1, 29]. Phenotypically, this study shows that both mutations are broadly similar, which is perhaps expected given that both mutations result in a similar functional perturbation.

Whilst these phenotypes are not entirely unique to NPC, as lipid storage, lysosomal expansion and autophagic vacuole accumalation are reminiscent of most lysosomal storage disorders (LSD), the additional presence of reduced lysosomal Ca^2+^ is a phenotype that has been reported in very few LSDs. Altered lysosomal Ca^2+^ homeostasis was first associated with loss of function of NPC1 [24] and, as we now show here, NPC2. The loss of NgBR function has previously been linked to the destabilisation and reduced expression of NPC2, leading to cholesterol storage [20]. The mechanism by which NPC2 may have reduced function in DHDDS is yet to be elucidated. Previous studies have demonstrated that an over-expression of NgBR results in increased ER localisation and stabilisation of NPC2 [20]. It is therefore tempting to speculate that levels of NgBR, or NgBR-NPC2 interaction, may be elevated in DHDDS as a compensatory mechanism, thus impacting NPC2 localisation. However, previous work has also shown that the overexpression of DHDDS or NgBR results in enhanced levels of its binding partner, indicating they act to stabilise each other, so a reduced expression of DHDDS can decrease NgBR expression [40]. Understanding how the expression and localisation of NgBR, and in turn NPC2, are affected by DHDDS mutations is an interesting avenue for future investigational and therapeutic studies.

Whilst an accumulation of cholesterol has been previously noted in NUS1 patient fibroblasts, nus1 knock-down zebrafish [21] and the fruit fly model [22], it has not been reported to the same extent for DHDDS. The work presented here supports a previous study that identified enlarged lysosomes containing electron-dense material as well as deposits of membranous whorls in a patient skin biopsy carrying an R211Q mutation [1]. Membranous whorls and electron dense material are commonplace in lysosomal storage patients [41, 42], with the whorl-like structures being attributed to gangliosides [43], such as the ganglioside GM1 we identified as elevated in DHDDS-patient lysosomes. We identified one previous report indicating the presence of elevated filipin staining in DHDDS fibroblasts from a heterozygous affected indivudal harbouring the R205Q mutation, and a minimal elevation in fibroblasts from a patient harbouring a heterozygous D95N mutation [3]. Our work corroborates the significantly elevated filipin in patients harbouring a R205Q mutation, and now confirms it is lysosomal. The comparitively minor filipin elevation present in the D95N is associated with a milder disease course, as suggested by the later onset of tonic-clonic seizures and ataxia. This correlation suggests lysosomal storage could be a biomarker of DHDDS, although this would need substantially more investigation.

The identification of NPC disease-like phenotypes, alongside the established interaction between NPC2 and DHDDS proteins [19], raises the possibility that therapeutics approved for NPC could also be beneficial to DHDDS, and indeed NUS1. Here we show that miglustat, the first approved therapeutic for NPC, is effective at reducing glycolipid storage and lysosomal swelling in DHDDS. Furthermore, it partially rescues the lysosomal Ca^2+^ phenotype, demonstrating that miglustat may provide a functional improvement. Over more than twenty years of NPC patient recorded use, miglustat has been shown to stabilise disease progression and even provide improvement in NPC patients, especially in those with milder disease progression where an increase in lifespan has now been reported [44, 45]. Very recently, two other small molecules were also approved for the treatment of NPC; arimoclomol [27] and N-acetyl-L-leucine [28]. Whilst the mechanism of action for both compounds is unclear, they both are reported to address to some degree, albeit less than miglustat, the pathogenic lipid storage and may be of interest for future DHDDS/NUS1 therapeutic studies.

## Acknowledgments

The authors would like to thank CureDHDDS for funding and for supporting this work in the Lloyd-Evans laboratory, HWE is supported by Action Medical Research (GN3018). We thank Drs Frances Elmslie and Eva Morava for providing the fibroblast cells. Confocal microscopy was performed at Cardiff University Bioimaging Hub Core Facility; we thank the team for their support.

---

**Supplementary Figure 1.**
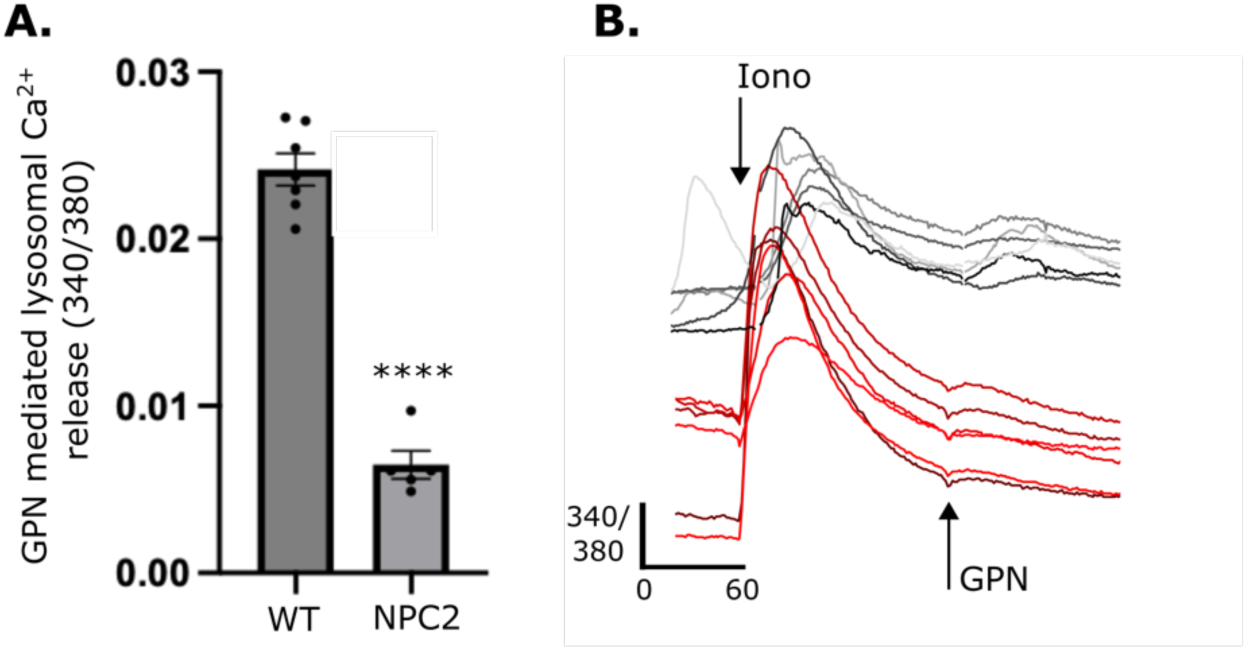
Lysosomal Ca^2+^ is reduced in NPC2 patient fibroblasts. **(A)** Lysosomal Ca^2+^ release was quantified from intracellular Ca^2+^ after GPN addition as detected by Fura 2,AM (ratiometric measurement at 340 nm and 380 nm, 340/380). Data is presented as the mean ± SEM, as determined by an unpaired t-test(****P ≤ 0.0001, n = 5 – 6). **(B)** Representative traces of NPC2 (shades of red) and apparently healthy control (shades of grey) patient fibroblasts loaded with Fura 2-AM were treated pharmacological agents, ionomycin (iono) and (GPN) to induce Ca^2+^ release.

## References

1. Galosi, S., et al., De novo DHDDS variants cause a neurodevelopmental and neurodegenerative disorder with myoclonus. Brain, 2022. 145(1): p. 208–223.

2. Hamdan, F.F., et al., High Rate of Recurrent De Novo Mutations in Developmental and Epileptic Encephalopathies. Am J Hum Genet, 2017. 101(5): p. 664–685.

3. Courage, C., et al., Progressive myoclonus epilepsies-Residual unsolved cases have marked genetic heterogeneity including dolichol-dependent protein glycosylation pathway genes. Am J Hum Genet, 2021. 108(4): p. 722–738.

4. Mehta, S. and V. Lal, DHDDS Mutation: A Rare Cause of Refractory Epilepsy and Hyperkinetic Movement Disorder. J Mov Disord, 2023. 16(1): p. 107–109.

5. Mousa, J., et al., Acetazolamide treatment in late onset CDG type 1 due to biallelic pathogenic DHDDS variants. Mol Genet Metab Rep, 2022. 32: p. 100901.

6. Williams, L.J., et al., DHDDS and NUS1: A Converging Pathway and Common Phenotype. Mov Disord Clin Pract, 2024. 11(1): p. 76–85.

7. Edani, B.H., et al., Structural elucidation of the cis-prenyltransferase NgBR/DHDDS complex reveals insights in regulation of protein glycosylation. Proc Natl Acad Sci U S A, 2020. 117(34): p. 20794–20802.

8. Adair, W.L., Jr., N. Cafmeyer, and R.K. Keller, Solubilization and characterization of the long chain prenyltransferase involved in dolichyl phosphate biosynthesis. J Biol Chem, 1984. 259(7): p. 4441–6.

9. Ericsson, J., et al., Characterization and distribution of cis-prenyl transferase participating in liver microsomal polyisoprenoid biosynthesis. Eur J Biochem, 1991. 202(3): p. 789–96.

10. Endo, S., et al., Identification of human dehydrodolichyl diphosphate synthase gene. Biochim Biophys Acta, 2003. 1625(3): p. 291–5.

11. Shridas, P., J.S. Rush, and C.J. Waechter, Identification and characterization of a cDNA encoding a long-chain cis-isoprenyltranferase involved in dolichyl monophosphate biosynthesis in the ER of brain cells. Biochem Biophys Res Commun, 2003. 312(4): p. 1349–56.

12. Park, E.J., et al., Mutation of Nogo-B receptor, a subunit of cis-prenyltransferase, causes a congenital disorder of glycosylation. Cell Metab, 2014. 20(3): p. 448–57.

13. Zelinger, L., et al., A missense mutation in DHDDS, encoding dehydrodolichyl diphosphate synthase, is associated with autosomal-recessive retinitis pigmentosa in Ashkenazi Jews. Am J Hum Genet, 2011. 88(2): p. 207–15.

14. Grünewald, S., G. Matthijs, and J. Jaeken, Congenital Disorders of Glycosylation: A Review. Pediatric Research, 2002. 52(5): p. 618–624.

15. Wen, R., B.L. Lam, and Z. Guan, Aberrant dolichol chain lengths as biomarkers for retinitis pigmentosa caused by impaired dolichol biosynthesis. Journal of lipid research, 2013. 54(12): p. 3516–3522.

16. Park, E.J., et al., Mutation of Nogo-B receptor, a subunit of cis-prenyltransferase, causes a congenital disorder of glycosylation. Cell metabolism, 2014. 20(3): p. 448–457.

17. Williams, L.J., et al., DHDDS and NUS1: A Converging Pathway and Common Phenotype. Movement Disorders Clinical Practice, 2024. 11(1): p. 76–85.

18. Vermeer, S., et al., Cerebellar ataxia and congenital disorder of glycosylation Ia (CDG-Ia) with normal routine CDG screening. Journal of neurology, 2007. 254: p. 1356–1358.

19. Kharel, Y., et al., In vivo interaction between the human dehydrodolichyl diphosphate synthase and the Niemann–Pick C2 protein revealed by a yeast two-hybrid system. Biochemical and biophysical research communications, 2004. 318(1): p. 198–203.

20. Harrison, K.D., et al., Nogo-B receptor stabilizes Niemann-Pick type C2 protein and regulates intracellular cholesterol trafficking. Cell metabolism, 2009. 10(3): p. 208–218.

21. Yu, S.-H., et al., Lysosomal cholesterol accumulation contributes to the movement phenotypes associated with NUS1 haploinsufficiency. Genetics in Medicine, 2021. 23(7): p. 1305–1314.

22. Xue, J., et al., Loss of Drosophila NUS1 results in cholesterol accumulation and Parkinson’s disease-related neurodegeneration. The FASEB Journal, 2022. 36(7): p. e22411.

23. Zervas, M., et al., Critical role for glycosphingolipids in Niemann-Pick disease type C. Current Biology, 2001. 11(16): p. 1283–1287.

24. Lloyd-Evans, E., et al., Niemann-Pick disease type C1 is a sphingosine storage disease that causes deregulation of lysosomal calcium. Nature medicine, 2008. 14(11): p. 1247–1255.

25. te Vruchte, D., et al., Accumulation of glycosphingolipids in Niemann-Pick C disease disrupts endosomal transport. Journal of Biological Chemistry, 2004. 279(25): p. 26167–26175.

26. Lachmann, R.H., et al., Treatment with miglustat reverses the lipid-trafficking defect in Niemann–Pick disease type C. Neurobiology of disease, 2004. 16(3): p. 654–658.

27. Mengel, E., et al., Efficacy and safety of arimoclomol in Niemann-Pick disease type C: results from a double-blind, randomised, placebo-controlled, multinational phase 2/3 trial of a novel treatment. Journal of inherited metabolic disease, 2021. 44(6): p. 1463–1480.

28. Bremova-Ertl, T., et al., Trial of N-Acetyl-l-Leucine in Niemann–Pick Disease Type C. New England journal of medicine, 2024. 390(5): p. 421–431.

29. Bar-El, M.L., et al., Structural basis of heterotetrameric assembly and disease mutations in the human cis-prenyltransferase complex. Nat Commun, 2020. 11(1): p. 5273.

30. Sugimoto, Y., et al., Accumulation of cholera toxin and GM1 ganglioside in the early endosome of Niemann–Pick C1-deficient cells. Proceedings of the National Academy of Sciences, 2001. 98(22): p. 12391–12396.

31. Harzer, K. and B. Kustermann-Kuhn, Quantified increases of cholesterol, total lipid and globotriaosylceramide in filipin-positive Niemann-Pick type C fibroblasts. Clinica Chimica Acta, 2001. 305(1): p. 65–73.

32. Vanier, M. and G. Millat, Niemann–Pick disease type C. Clinical genetics, 2003. 64(4): p. 269–281.

33. Choudhury, A., et al., Rab proteins mediate Golgi transport of caveola-internalized glycosphingolipids and correct lipid trafficking in Niemann-Pick C cells. Journal of Clinical Investigation, 2002. 109(12): p. 1541–1550.

34. Nielsen, G.K., et al., Protein Replacement Therapy Partially Corrects the Cholesterol-Storage Phenotype in a Mouse Model of Niemann-Pick Type C2 Disease. PLOS ONE, 2011. 6(11): p. e27287.

35. Naureckiene, S., et al., Identification of HE1 as the Second Gene of Niemann-Pick C Disease. Science, 2000. 290(5500): p. 2298–2301.

36. Pagano, R.E., Endocytic trafficking of glycosphingolipids in sphingolipid storage diseases. Philos Trans R Soc Lond B Biol Sci, 2003. 358(1433): p. 885–91.

37. te Vruchte, D., et al., Relative acidic compartment volume as a lysosomal storage disorder–associated biomarker. The Journal of Clinical Investigation, 2014. 124(3): p. 1320–1328.

38. Morgan, A.J., et al., Does lysosomal rupture evoke Ca2+ release? A question of pores and stores. Cell Calcium, 2020. 86: p. 102139.

39. Yuan, Y., et al., The lysosomotrope GPN mobilises Ca2+ from acidic organelles. Journal of Cell Science, 2021. 134(6).

40. Harrison, K.D., et al., Nogo-B receptor is necessary for cellular dolichol biosynthesis and protein N-glycosylation. EMBO J, 2011. 30(12): p. 2490–500.

41. Barral, D.C., et al., Current methods to analyze lysosome morphology, positioning, motility and function. Traffic, 2022. 23(5): p. 238–269.

42. Vogler, C., et al., Electron microscopy in the diagnosis of lysosomal storage diseases. Am J Med Genet Suppl, 1987. 3: p. 243–55.

43. Ferreira, C.R. and W.A. Gahl, Lysosomal storage diseases. Transl Sci Rare Dis, 2017. 2(1-2): p. 1–71.

44. Pineda, M., M. Walterfang, and M.C. Patterson, Miglustat in Niemann-Pick disease type C patients: a review. Orphanet J Rare Dis, 2018. 13(1): p. 140.

45. Patterson, M.C., et al., Miglustat for treatment of Niemann-Pick C disease: a randomised controlled study. Lancet Neurol, 2007. 6(9): p. 765–72.

